# «Involvement of Pf-like phages in resistance to phage infection in clinical isolates of *P. aeruginosa* from cystic fibrosis patients*»*

**DOI:** 10.1101/2024.05.03.592413

**Authors:** Lucía Blasco, Antonio Barrio-Pujante, Carla López-Causape, Lucía Armán, Inés Bleriot, Olga Pacios, Laura Fernández-García, Concha Ortiz-Cartagena, Clara Ibarguren-Quiles, Rafael Cantón, Antonio Oliver, María Tomás

## Abstract

*Pseudomonas aeruginosa* is a bacterial pathogen that is a major cause of lung infections in cystic fibrosis (CF) and other patients. Isolates of *P. aeruginosa* from CF patients commonly carry filamentous phages (Pf phages), a type of temperate phage known to be related to biofilm production and antibiotic sequestration. In this study, 12 new Pf-like phages were identified in a collection of clinical isolates of *P. aeruginosa* from CF patients. Analysis of the phage genomes revealed different anti-phage defence systems, described here for first time in these types of phages. Finally, relationships between resistance to phage infection and the presence of Pf-like phages and also between each defence system and resistance were observed.

**IMPORTANCE:** Bacteria harbour a wide range of defence mechanisms to avoid phage infections. These mechanisms hamper the application of phage therapy because they can lead to the rapid acquisition of phage resistance. Temperate phages, including the filamentous phages, carry genes encoding virulence factors and also anti-phage defence mechanisms, as their survival depends on the host survival. In this study, we identified 12 new Pf-like phages encoding several different anti-phage defence mechanisms, some observed for the first time in this type of phage. A relationship between these phages and resistance to phage infection was also observed. The study findings are important as they provide information about newly discovered filamentous phages and their proteomes and also about the role of these phages in resistance to phage infections. Studying the genome of clinical isolates carrying these phages could help to improve phage therapy by targeting these phages or its genes.

## INTRODUCTION

The interaction between bacteria and the viruses that infect them, phages, is an evolutionary driving force. Both organisms coevolve in an “arms race”, with many different immune mechanisms developed in bacteria and counterpart mechanisms developed in phages.

The filamentous phages belong to the order *Tubulavirales* (1). These phages are uncommon both in morphology and life cycle. They are present in the host genome as prophages and when assembled, they exit the cell by extrusion without lysing the bacteria, causing chronic infections (7). They have a helical structure composed by the major coat protein, which surrounds circular, positive-sense, single stranded DNA. Many phage species integrate their genome in the host genome, but others such as episomal phages are non-integrative (1, 2).

The filamentous phages are widely distributed in the multiresistant pathogen *Pseudomonas aeruginosa*. This important pathogen, designated “high risk” by the World Health Organization (WHO) in 2017 (3), can cause severe infections in hospitals and is responsible for chronic infections in the respiratory tract, wounds and burns (4). *P. aeruginosa* is also closely associated with infections in cystic fibrosis (CF) patients, at least partly because of its ability to form dense biofilms, which favours the development and chronicity of the infection (4). The filamentous phages are found within these biofilms as crystal liquid structures called tactoids, which can enhance phage tolerance to antibiotics by forming an adsorptive diffusion barrier (5). The filamentous phages identified in *P. aeruginosa* are designated Pf-like phages, of which seven types have been described to date. It is estimated that 50 to 60 % of *P. aeruginosa* isolates are lysogenized by a Pf-like phage. The high prevalence of these phages in *P. aeruginosa* is related to their role in pathogenesis, virulence and immune system evasion (7).

Pf4 is a filamentous phage that infects *P. aeruginosa* and whose genomic structure is typical of the integrative *P. aeruginosa* filamentous phages. The genome of Pf4 and other Pf phages is divided into a conserved part, the core genome, and a non-conserved part, the accessory genome. The core genome is the part of the genome required for completion of a replication cycle in Gram-negative hosts, and it comprises genes related to structure, replication, assembly and secretion (6). By contrast, the accessory genome is a variable part of the genome which contains many genes of unknown function, as well as toxin genes, whose function is related to interaction with the host and involves virulence factors or toxin-antitoxin systems (6).

The anti-phage defence mechanisms, which are considered the “prokaryotic immune system”, are encoded in mobile elements in the bacterial genome, such as defence islands and prophages (7). These defence systems are frequently organized in gene clusters (8). As these systems are present in mobile genetic elements, they are usually acquired by bacteria through horizontal gene transfer, promoting environmental adaptation of the bacterial communities (9). Different anti-phage defence systems are encoded in clusters that are used as immune strategies in bacteria and include the following: a) Adsorption resistance, which is the first barrier to infection. Bacteria can evade adsorption by hiding the receptors with extracellular polymers or by mutations in the receptor gene. These mutations involve loss of receptors or structural changes in the receptors; b) Prevention of host takeover, which occurs after phage adsorption and prevents irreversible takeover of the host metabolism. This can be achieved by the Restriction-Modification (RM) systems, which are conformed by a restriction endonuclease and a methyltransferase. RM systems act by restricting the phage genome and methylation of the host genome thus protecting it from the endonuclease action. The Clustered Regularly Interspaced Short Palindromic Repeats (CRISPR)-associated proteins (CRISPR-Cas) form an adaptative immune system characterized by the acquisition of small fragments of foreign DNA, known as spacers, between the CRISPR locus repeats. The spacers are used to recognize exogenous nucleic acids, which will be degraded by the Cas endonuclease. Superinfection exclusion (Sie), a defence system developed by prophages or plasmids present in the host, blocks the uptake of phage nucleic acid into the cytoplasm. c) Abortive infection systems (Abi systems), which are different systems that inhibit the infection at any of the stages of DNA replication, translation or transduction, so that phages are unable to infect the bacteria, and the bacteria die or become persistent; and d) the Toxin-Antitoxin (TA) system, which acts by reducing the bacterial metabolism and thus inhibiting phage replication under stress conditions (9–13).

In this study, we examined the prevalence of Pf-like phages in 75 clinical isolates of *P. aeruginosa* from 25 chronic CF patients; we also examined the relationship between the prophages and the host resistance to phage infection.

## RESULTS

### Prevalence of Pf-like phages in clinical isolates of *P. aeruginosa* from CF patients

The genome of 75 clinical isolates of *P. aeruginosa* from 25 chronic CF patients (3 isolates per patient) were analysed to search for complete genomes of filamentous phages. The PHASTER search reported the presence of complete genomes of 42 filamentous phage distributed in 39 isolates and also 36 isolates without filamentous phage genome (Table 1). The presence of filamentous prophages in all isolates derived from one patient was variable; thus, in 40% of the patients all isolates carried a filamentous phage; in 12% of patients the filamentous phage was present in two isolates and in the other 12% only one isolate carried a filamentous phage. Finally in 36% of the patients, none of the isolates contained a filamentous phage in the genome (Table 1). The isolates from a patient in which no filamentous phage was found belonged to a different sequence type (ST) than the other isolates from the same patient, which carried filamentous phage (except in the case of patient 24.)

**Table 1.** Clinical strains *of P. aeruginosa* recovered from CF patients, showing the isolates grouped by patient, the ST and the filamentous phage found in each isolate. The lytic phages used in the study are also shown.

| Patient | *P. aeruginosa* CF isolate | ST | Pf-like isolate |
| --- | --- | --- | --- |
| 01 | 01-0440 | 1089 | Pf01-0440 |
|  | 01-5978 | 1089 | - |
|  | 01-7071 | 1089 | Pf01-7071 |
| 02 | 02-5135 | 312 | Pf02-5135a;Pf02-5135b |
|  | 02-5867 | 312 | Pf02-5867a;Pf02-5867b |
|  | 02-6433 | 312 | Pf02-6433a;Pf02-6433b |
| 03 | 03-0062 | 285 | Pf03-0062 |
|  | 03-5302 | 285 | Pf03P. -5302 |
|  | 03-5453 | 285 | Pf03-5453 |
| 04 | 04-5265 | 274 | Pf04-5265 |
|  | 04-7991 | 274 | Pf04-7991 |
|  | 04-8869 | 274 | Pf04-8869 |
| 05 | 05-2269 | NEW1 | - |
|  | 05-2840 | NEW1 | - |
|  | 05-4672 | 360 | Pf05-4672 |
| 06 | 06-6855 | 242 | - |
|  | 06-7209 | 242 | - |
|  | 06-9800 | 242 | - |
| 07 | 07-1155 | 279 | - |
|  | 07-5966 | 279 | - |
|  | 07-8998 | 279 | - |
| 08 | 08-1318 | NEW2 | Pf08-1318 |
|  | 08-4371 | NEW2 | Pf08-4371 |
|  | 08-5924 | NEW2 | Pf08-5924 |
| 09 | 09-0786 | 1109 | Pf09-0786 |
|  | 09-3048 | 1109 | Pf09-3048 |
|  | 09-9593 | 1109 | Pf09-9593 |
| 10 | 10-6443 | 360 | Pf10-6443 |
|  | 10-6518 | 360 | Pf10-6518 |
|  | 10-7858 | 360 | Pf10-7858 |
| 11 | 11-4349 | 198 | - |
|  | 11-7257 | 2475 | Pf11-7257 |
|  | 11-8664 | 2475 | Pf11-8664 |
| 12 | 12-0969 | 277 | - |
|  | 12-2742 | 277 | - |
|  | 12-2760 | 277 | - |
| 13 | 13-0154 | 1123 | Pf13-0154 |
|  | 13-1387 | 1123 | Pf13-1387 |
|  | 13-2748 | 1123 | Pf13-2748 |
| 14 | 14-4114 | 319 | - |
|  | 14-4688 | 319 | - |
|  | 14-5818 | 319 | - |
| 15 | 15-4963 | 412 | - |
|  | 15-7676 | 412 | - |
|  | 15-8860 | 412 | - |
| 16 | 16-0109 | 252 | - |
|  | 16-2856 | 252 | - |
|  | 16-4264 | 408 | Pf16-4264 |
| 17 | 17-0755 | 274 | - |
|  | 17-3115 | 274 | - |
|  | 17-8321 | 274 | - |
| 18 | 18-6285 | 312 | Pf18-6285 |
|  | 18-7126 | 312 | Pf18-7126 |
|  | 18-9439 | 312 | Pf18-9439 |
| 19 | 19-0943 | 1092 | Pf19-0943 |
|  | 19-6618 | 1092 | Pf19-6618 |
|  | 19-9746 | 1092 | Pf19-9746 |
| 20 | 20-0447 | 1072 | Pf20-0447 |
|  | 20-3695 | 1072 | Pf20-3695 |
|  | 20-6028 | 1072 | Pf20-6028 |
| 21 | 21-2955 | 198 | - |
|  | 21-4234 | 198 | - |
|  | 21-9889 | 198 | - |
| 22 | 22-5179 | NEW3 | - |
|  | 22-5546 | 2101 | - |
|  | 22-5835 | 1134 | - |
| 23 | 23-2344 | 701 | - |
|  | 23-6966 | CC701 | - |
|  | 23-9557 | 701 | - |
| 24 | 24-0501 | 274 | Pf24-0501 |
|  | 24-1092 | 274 | Pf24-1092 |
|  | 24-7416 | 274 | - |
| 25 | 25-6546 | 1072 | Pf25-6546 |
|  | 25-7986 | 235 | - |
|  | 25-9260 | 1613 | - |
| ***P. aeruginosa* reference strain** |  | **Origin** |  |
| PA01 |  | Present study |  |
| PA14 |  | Present study |  |
| **Lytic Phages** |  | **Origin** |  |
| φDCL-PA6 |  | Contaminated river water (Contreras research group, UNAM) (47) |  |
| φDCL-PA6α |  | Contaminated river water (Contreras research group, UNAM)(47) |  |
| PAC8 |  | Contaminated river water (Contreras research group, UNAM)(47) |  |
| PAC2 |  | Compost (This group) |  |

As the filamentous phages present in *P. aeruginosa* are known as Pf phages, the phages identified in this study will be named in the same way as Pf-like phages in general, with the number of the isolate added, as appropriate (Table1).

### Phylogenetic analysis of the Pf-like phages

The Pf-like phage genomes identified were phylogenetically analysed, and the maximum likelihood tree revealed a high degree of homology (Fig. 1A). Although the genomes were grouped by patient, some clades showed a high degree of similarity between phages from different patients. The first cluster was constituted by the Pf-like genomes located in patients 01, 04 and 24. A second cluster consisted of the Pf-like phages identified in patients 02 and 18, and a third cluster was formed by the phages isolated from patients 20 and 25. A fourth cluster included the phages isolated from patients 10 and 05 (Fig. 1A, 1B). A Brig BLAST and an ANI study comparing the phage genome sequences of each tree clade revealed a high level of homology, of between 99.92% and 100%, sowere assumed to be of the same phage (Fig. 1B). Based on these results, a total of 12 Pf-like phages were identified. The genome sequences are included in Bioproject PRJNA1082103 (Table 2).

**Figure 1.**
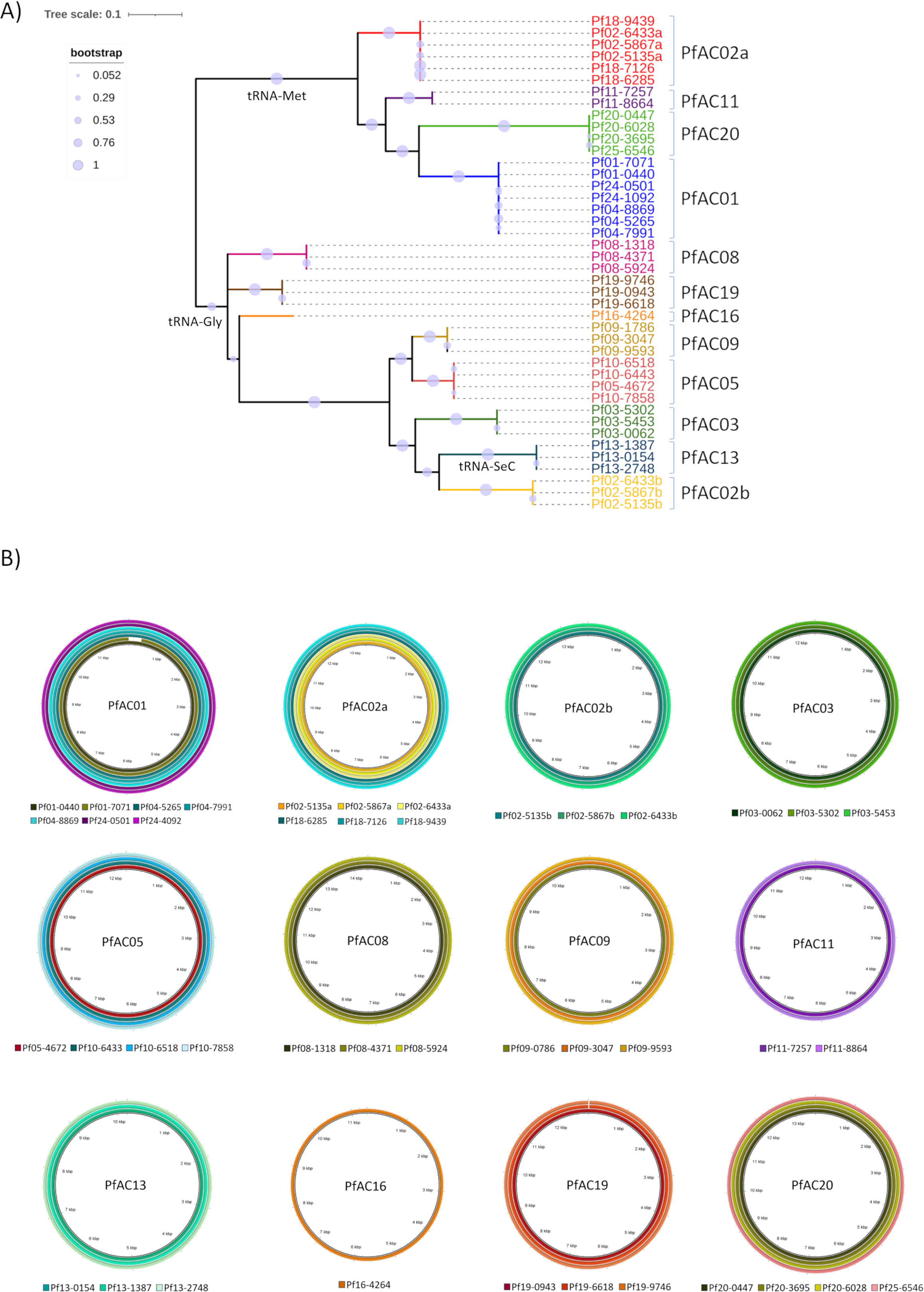
Genomic analysis of the filamentous phage genomes. (A) Phylogenetic analysis by Maximum likelihood, with the tree showing the groups that corresponded to the final Pf-like phages identified and two major groups corresponding to the insertion sites. (B) Brig homology analysis of the filamentous phage genome groups.

**Table 2.**
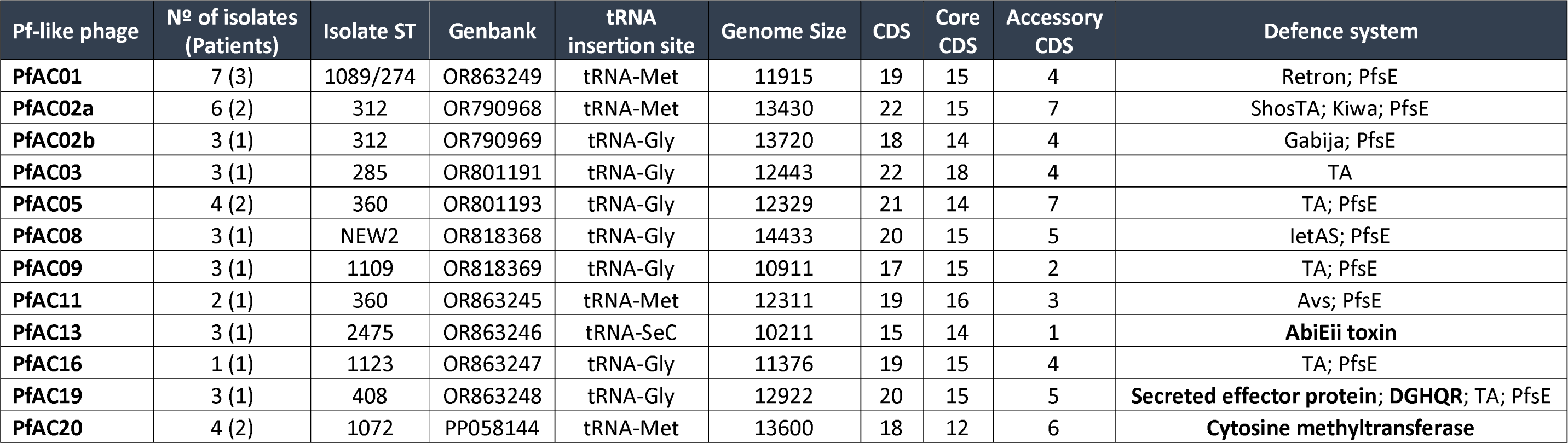
12 Pf-like phages identified in this study, showing the genome size, Genbank code, ST related to each phage, insertion site for each Pf-like phage, CDS number, Core Genome CDS, Accessory Genome CDS and Defence System found in each Pf-like phage.

The maximum likelihood tree was divided into two major groups corresponding to the bacterial attachment site (attB) of the prophages. One group was constituted by the Pf-like genomes that use the tRNA-Met as attB and the other group included those with the tRNA-Gly and tRNA-Sec attB sites (Table 2; Fig. 1A). Four of 5 phages with a tRNA-Met attB were present in isolates from different patients, while the phages with the tRNA-Gly and tRNA-Sec were only present in the isolates from one patient, with the exception of phage PfAC05, which was present in two patients (Fig. 1A).

### Genomic analysis of the filamentous Pf-like prophages

Analysis of the genomes showed that the 12 Pf-like phages identified were integrated in different tRNA sites, so that 45% were integrated in a tRNA-Met, 47.5% in a tRNA-Gly and 7.5% in a tRNA-Sec.

The genomic analysis revealed that all of the filamentous phages were of between 10kb and 14Kb in size and had between 15 and 22 CDS.

Annotation of the genes from the 12 Pf-like phages showed that the genome structure comprised a core genome composed by 12 to 18 CDS, flanked by 1 to 7 CDS corresponding to the accessory genome (Fig. 2A). The core genome was composed by genes encoding structural and capsid proteins as well as proteins involved in replication and flanked by two integration proteins, an integrase and an excisionase. The core genome was conserved across all of the Pf-like phages identified (Fig. 2A). The accessory genome flanked the core genome and was composed by moron genes, mainly related to anti-phage defence, but also genes encoding for hypothetical proteins as well as ATP-binding proteins and Arc family DNA-binding proteins.

**Figure 2.**
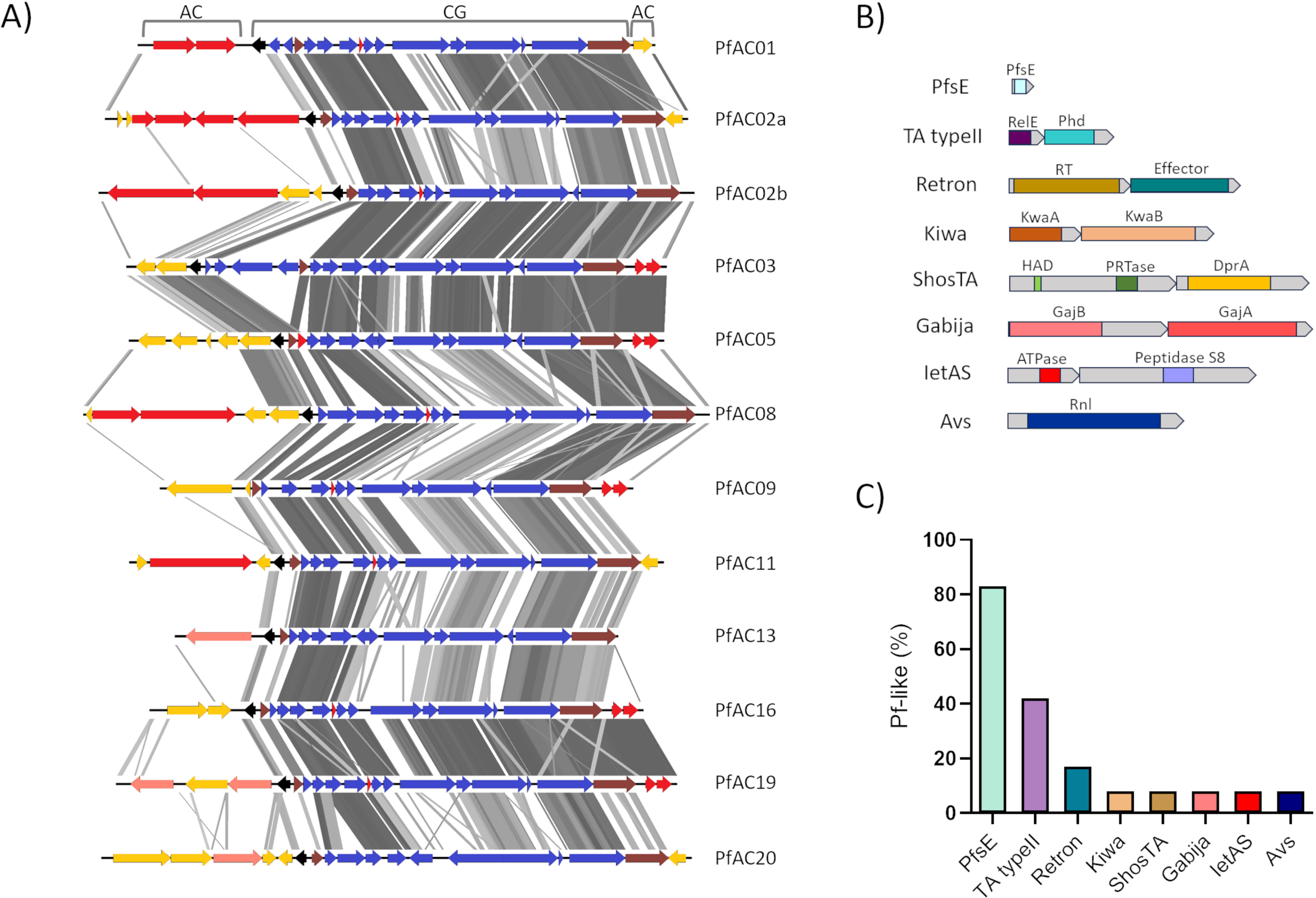
Pf-like phage genome and anti-phage defence systems. (a) Protein annotation and homology of the 12 Pf-like phages. The Core Genome (CG), Accessory Genome (AG), integrase and excisionase (brown), excisionase negative regulator (black), defence systems (red), hypothetical genes in the accessory genome (yellow) and genes of the core genome (blue) are shown. (B) Protein components of the defence systems. (C) Prevalence of each defen**c**e system in the 12 Pf-like phages identified.

The genes belonging to a complete defence system constituted 36% of the accessory genome, and 7% were isolated proteins from incomplete defence systems. All of the Pf-like phages identified, carried two complete defence systems, except phage PfAC02a, which carried 3, systems, and PfAC13 and PfAC20, which only had incomplete systems (Fig. 2A; Table2).

The following 8 complete defence systems were detected: TA typeII, Retron, Kiwa, ShosTA, Gabija, IetSA, PfsE and Avs (Fig. 2A, 2B, 2C; Table 2). The PfsE gene, unlike the other defence systems, was located in the core genome. This gene was present in 10 of the 12 Pf-like phages (83%). The second most common system was the TA TypeII system, which was present in 5 Pf-like phages (42%). Annotation showed that this system corresponded to a cluster composed by two contiguous genes, a RelE family toxin and a Phd family antitoxin. The retron system was present in two Pf-like phages (17%) and according to the protein annotation it was a cluster constituted by a retrotranscriptase and a retron effector protein. Both Kiwa and ShosTA systems were present in the PfAC02a Pf-like (8%). Kiwa was a gene cluster composed by kwaA and kwaB genes, identified by PADLOC. ShosTA was also a cluster constituted by two genes, a DNA-binding protein (DprA-like) and a phosphoribosyl transferase (PRTase). Finally, both IetAS and Avs were present in one Pf-like phage (8%). The annotation revealed that the IetAS system was composed by a peptidase S8 (IetS) and a putative ATPase (IetA), while the Avs was constituted by an NLR ATPase identified by HHPred.

Each of the incomplete systems were represented in 8% of the Pf-like phages. One gene coding for the putative AbiEii toxin was found in phage PfAC13; two genes from incomplete systems, a Secreted effector protein and DGHQR, were found in phage PfAC19, and finally a DNA cytosin methyltransferase was present in phage PfAC20.

### Phage resistance pattern

All 75 clinical isolates were infected with 4 lytic *P. aeruginosa* phages in order to study the relationship between phage resistance and the presence of defence systems in the accessory genome of the Pf-like phages.

The infection study was conducted by spot testing and broth infection curves. From a total of 300 phage-bacteria interactions, 223 (74.33%) were resistant and 77 (25.66%) sensitive (Fig. 3A). Of the interactions between the 4 lytic phages and 39 Pf-like carrying isolates, 124 resulted in resistant interactions that were significantly higher than the 99 resistant interactions with the non-Pf-like phages. By contrast, the sensitive interactions were significantly higher in the non-Pf-like carriers (Fig. 3A, 3B). Finally, although both carrier and non-carrier Pf-like isolates were mainly resistant to phage infection, the probability of an isolate being resistant when it carried a Pf-like phage was 15.6% higher than when it did not carry any Pf-like phage (Fig. 3C).

**Figure 3.**
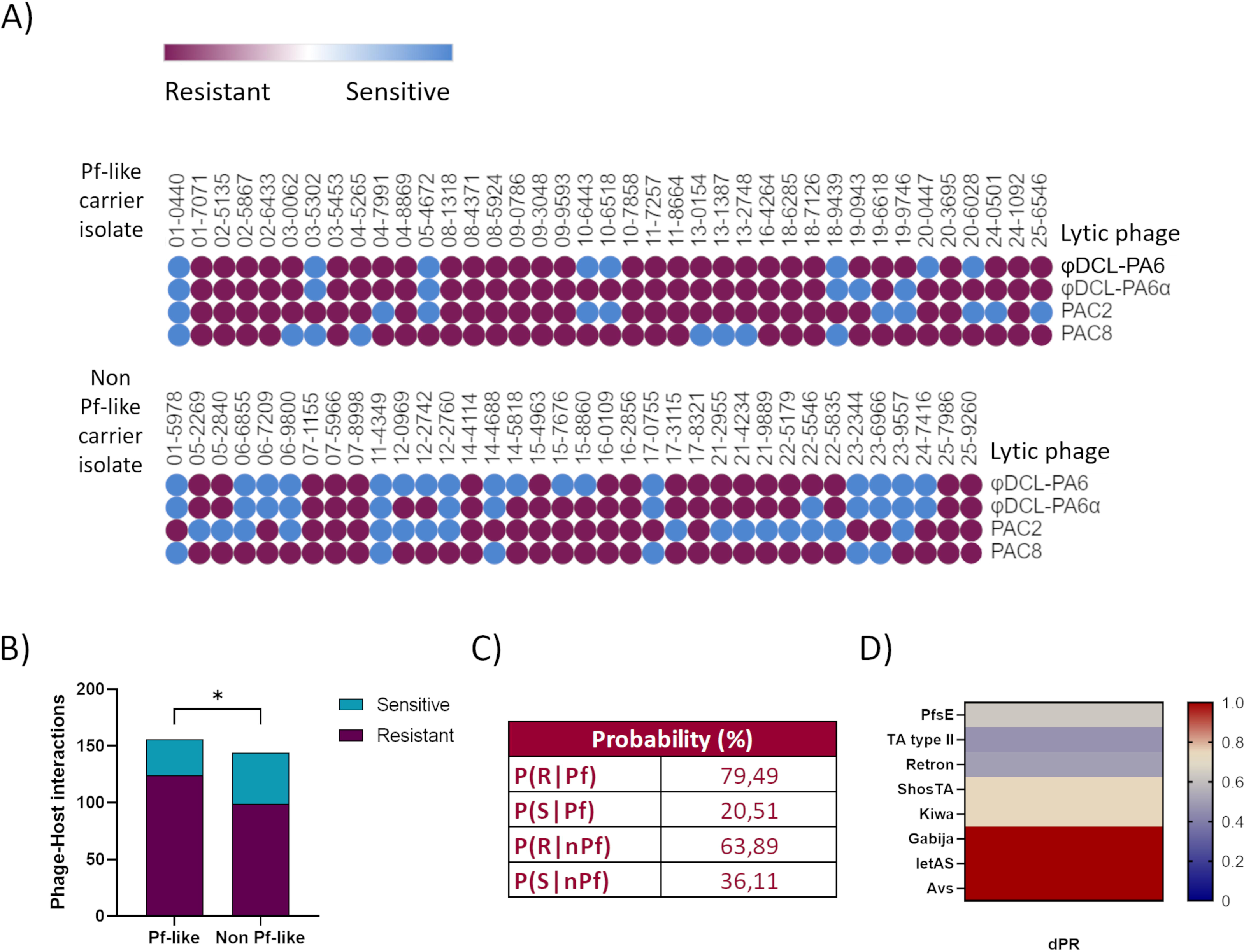
Relationship between the Pf-like phage and resistance to phage infection. (A) Resistance pattern for each isolate (Pf-like carrier and non-Pf-like carrier) challenged with 4 lytic phages; the graph represents the results obtained in both the spot test and infection curves. (B) Resistant and Sensitive interactions between the *P. aeruginosa* identified as carriers and non-carriers of Pf-like phages. (C) Probability that the *P. aeruginosa* isolates identified as carriers and non-carriers of Pf-like phages are resistant to phage infection. (D) Differential probability (dPR) of each anti-phage defence system carried in the Pf-like genomes. Values of dPR close to 1 indicate overrepresentation of phage resistance, while negative values indicate overrepresentation of sensitivity to phage infection.

The relationship between resistance and each defence system in Pf-like phage carrier isolate, was calculated by differential probability (Fig. 3D). All of the systems yielded values of dPR>0, indicating that they are probably related to phage resistance. Gabija, IetAS and Avs yielded the highest value of dPR=1, which indicates that these systems were directly related to resistance. The probability that ShosTA and Kiwa were involved in resistance was equal because these were present in the same phage and it was not possible to differentiate the individual activity of each. By contrast, the presence of Gabija with ShosTA and Kiwa in the isolates from patient 02 increased the differential probability to 1, indicating a synergic effect. The PfsE system co-occurred with all other systems, but no synergistic effect was observed, as the differential probability was different for each phage.

## DISCUSSION

The coexistence of bacteria and other microorganisms in the environment leads to competitive, collaborative and predatory interactions. Bacteria have developed different systems to manage these interactions, e.g. secretion systems and phage tail-like bacteriocins, which are involved in competition with other bacteria, and defence systems, which are used to evade phage infections (9, 10, 14). The diversity of systems that make up the bacterial defence arsenal is largely driven by the “arms race” between bacteria and phages and is, in turn, enhanced by horizontal gene transfer of mobile genetic elements such as defence islands and temperate phages (15, 16). During the lysogenic phase, temperate phages maintain a symbiotic relationship with their hosts, and the fitness of both is intimately linked. The presence of virulence and defence genes in the accessory genome of the temperate phages increases the survival of the both the host and the phage itself (13, 15, 17).

In this study genomic analysis of 75 clinical isolates recovered from 25 CF patients led to the identification of 42 filamentous phage genomes encompassed in the Pf-like type phages. Phylogenetic and homology studies showed that the 42 Pf-like genomes corresponded to 12 different Pf-like phages, described here for first time. The genomes of these prophages were found disrupting attB tRNA-Met or tRNA-Gly in the same proportion and tRNA-Sec in a lower proportion. This distribution was previously reported by Fiedoruk *et al.* (2020) (4, 18). However, we observed a relationship between the tRNA attachment site and the homology between the phages, represented in the phylogenetic tree as 2 major clades (Fig. 1A).

The type of Pf-like phage is closely related to the ST of the isolate, which explains the absence of Pf-like phages in isolates from the same patient but belonging to different STs. The acquisition of antibiotic resistance and its maintenance in clinical *P. aeruginosa* clones was related to exposure to certain antibiotics and the acquisition of resistance genes by horizontal transfer or mutations. A relationship between continuous exposure of the *P. aeruginosa* isolates to antibiotics and the prevalence of the Pf-like phages due to its role sequestering the antibiotics in the biofilm was also described. Thus, the observed relationship between the presence of a Pf-like and a ST may depend on the antibiotics used to treat the CF patient (19–21). This relationship may be of interest for typing of the isolates of *P. aeruginosa*, but further studies are needed to confirm its existence.

Annotation of the Pf-like genome and assignation of gene function revealed a canonical organization of the genes in a core genome flanked by an accessory genome. The core genome consists of a variable number of genes, between 10 and 16 genes, with different functions: morphogenesis, assembly, DNA replication, integration and excision (4). The organization of the core genome of the Pf-like phages identified was similar to that of the Pf4 phage, which is widely used as a Pf-like model. The following common genes were identified: C repressor gene pf4r (PA0715); excisionase XisF4 (PA0716); single-stranded DNA binding protein (PA0720); coaB, major coat protein (PA0723); coaA, minor coat protein (PA0724); Zot domain protein (PA0726); replication initiation protein (PA0727) and integrase intF (PA0728) (4, 18).

The 12 Pf-like phages identified were very similar in the core genome region but differed in the genes carried in the accessory genome (Fig. 3A). The accessory genes of the Pf-like phages identified were analysed, and a function was assigned to approximately half of the genes, depending on the phage. The number of genes varied between 1 and 6, as previously described for other Pf-like phages (4, 18). The accessory genome of Pf-like phages, shared by other prophage families, was described as a group of genes that are not essential for the virus but with functions that benefit the host and improve its survival (15). The beneficial genes present in some prophages include anti-phage defence systems. To date, only TA and PfsE defence systems have previously been described in a filamentous phage (22). Eight defence systems were identified in the genomes of different Pf-like in this study, and the presence of these Pf-like phages was also found to confer greater resistance to phage infection than when they are not present. As previously described in other families of temperate phages, the presence of these Pf-like phages can enhance host survival, protecting bacteria from lytic phages via different mechanisms including inhibition of DNA translocation, premature transcription termination and abortive infection (15). The anti-phage defence systems identified in the Pf-like phages in this study were involved in different defence mechanisms. From the different defence strategies, we found systems representative of adsorption resistance (PfsE), Abi (Retron, Kiwa, Gabija, Avs) and TA (TA typeII, ShosTA, IetAS) (7, 23).

The probability of the occurrence of a defence system was, in all cases, related to phage resistance in the clinical isolate. However, it was observed that highly prevalent anti-phage defence systems were less likely to be related to resistance. This is consequence of the “arms race” between bacteria and phages, as more prevalent systems will be involved in a greater number of interactions with lytic phages, which will therefore be more likely to develop counterdefensive mechanisms (6).

Among the 12 Pf-like phages identified, the most common defence system was the pfsE gene (observed in 83% of the Pf-like phages). This was the only gene located in the core genome, which explains its high prevalence as this region is conserved in the Pf-like phages (Fig. 2A, 2B, 2C). The PfsE protein, first identified in the Pf4 *P. aeruginosa* filamentous phage, provides resistance to adsorption by suppressing extension of the pilus type 4 (which acts as a receptor for many lytic phages) via binding to PilC (24). This protein has also been identified as an inhibitor of the Pseudomonas quinolone signal (PQS) quorum sensing (25). The high prevalence of PfsE contrasts with the lower probability of involvement in phage resistance than other less frequent systems (Fig. 3D), which may be a result of its role in the suppression of the pilus as phage receptor, which would only be useful for inhibiting infection by phages using this receptor.

Of the defence systems present in the accessory genome, the TA system was most prevalent (43%) and was less likely to be involved in resistance than PfsE (Fig. 2B, 2C). The TA system identified belongs to the type II TA systems, which is composed by a genetic module encoding a toxin-antitoxin system, where the antitoxin protein blocks the toxin protein by protein-protein interactions (12, 22). Although the type II TA systems are involved in inhibiting the central cellular roles such as DNA replication and translation, they have a primarily biological role in inhibiting phage infection (5, 39). As in this study, a type II TA system was presentin the Pf4 filamentous phage, in which the toxin protein belongs to the ParE family and the antitoxin to the PhD family (22, 26). As with PfsE, the high frequency of this system in the bacterial and filamentous phage genomes has favoured the development of anti-TA mechanisms by lytic phages (10).

A retron system was present in 17 % of the Pf-like phages analysed and was estimated to have high probability of being involved in phage resistance (Fig. 2B, 2C; Fig. 3D). Retrons encode a specialized reverse transcriptase and a unique chimeric single-stranded DNA/RNA molecule. Although their existence has been known for more than 30 years, it was not until 2020 that their role in phage defence was determined (27, 28). The relationship between retrons and anti-phage defence was also observed in the present study (Fig. 3D).

The other five defence systems identified, Gabija, Kiwa, ShosTA, IetAS and Avs, were present in 8 % of the Pf-like phages, but their role in the resistance against phage infection was variable, with a direct relationship for Gabija, IetAS and Avs (dPR=1) (Fig. 3D). Gabija was recently partly identified as a nucleotide-sensing endonuclease (15). The Gabija system is composed by two genes, *gajA* and *gajB*, where *gajA* encodes a specific DNA nicking endonuclease and *gajB* encodes a helicase. The GajA endonuclease is activated by the depletion of NTP and dNTP when transcription of phage DNA occurs. It has been speculated that that, as a helicase, GajB may interact with GajA and somehow stimulate the binding, cleavage and/or turnover of GajA (29).

The IetAS system is also directly related to the phage resistance of the strain, but the mechanism of action remains unknown (15).

In the case of Kiwa and ShosTA, the value of the differential probability of involvement in phage defence (0.6) indicates overrepresentation of these systems in the resistant isolates. However, both systems were found in the same phage and their individual role in defence was not determined. The ShosTA system is a TA system composed by two proteins, a DprA-like protein as an antitoxin and a phosphoribosyl transferase (PRTase) as a toxin (30). The defence mode of action has been proposed to consist of detection of the phage by the DprA-like protein and activation of the PRTase, triggering the mechanisms for Abi (15). The Kiwa system was characterized as an Abi defence system constituted by two proteins, KiwaA and KiwaB. KiwaA detects the inhibition of the RNA polymerases by the lytic phage proteins and activates KiwaB, which reduces the phage DNA replication in a RecBCD-dependent manner (7).

The Avs system has previously been related to phage resistance and proposed to provide specific sensors for conserved structural features in phage proteins, such as the large terminase subunit and phage portal protein. It has been suggested that Avs tetramerize and activate an effector-mediated Abi-like response (7).

The great diversity of anti-phage defence systems found in the Pf-like phages identified may be a result of the co-existence of bacterial and phages in CF mucus, as recently demonstrated in a study conducted in the fish pathogen *Flavobacterium columnare*, in which co-existence with a predatory phage was involved in the development of phage resistance, in particular by the acquisition of CRISPR-Cas immunity (31, 32). Both lytic and lysogenic phages are ubiquitous in the body and therefore in the lungs of people suffering from CF. The mucus in the lungs of CF patients is hyper-concentrated and has a unique structure that favours bacterial colonization and, as also occurs in gut mucosa, the phages bind to the mucus. The mucus creates spatial refuges that favour the coexistence between phages and bacteria, which can explain the co-evolution of both (32–34).

To our knowledge this is the first time in which all of these defence systems have been identified in Pf-like phages and related to a high degree of phage resistance in Pf-like carrying isolates of *P. aeruginosa*. Pf-like phages have been linked to virulence traits and confer a competitive and survival advantage to the bacteria that possess them. The presence of defence systems in all of the isolates under study here suggests that the action of these phages favours survival of the *P. aeruginosa* isolates recovered from CF patients, both by increasing its virulence and by providing increased resistance to phage infection. Study of the presence of the Pf-like phages and the presence of defence systems in the genomes may, together with the relationship with ST, be of interest to improve the phage therapy by facilitating selection of appropriate lytic phages.

## MATERIAL AND METHODS

### Bacterial and lytic phage strains

Seventy-five *P. aeruginosa* clinical isolates were recovered from 25 CF patients (3 isolates per patient) (Table 1). The isolates, belonging to 26 STs from a collection of *P. aeruginosa* isolates from CF patients, were provided by the research group led by A. Oliver (Sons Espases Hospital, Palma de Mallorca, Spain) (35). *P. aeruginosa* PA01 and PA14 were used to propagate the lytic phages.Four *P. aeruginosa* lytic phages were used in the study (Table 1)

The *P. aeruginosa* isolates were cultured in LB (0.5% yeast extract; 0.5% NaCl; 1% tryptone), and agar 2% was added when necessary.

### Genome sequencing of the *P. aeruginosa* clinical isolates and filamentous phage genome identification and annotation

Next Generation Sequencing (NGS) of the isolates was performed in a previous study, with the MiSeq sequencing system (Illumina platform). The sequences were assembled using the Newbler Roche assembler and Velvet (Velvet v1.2.101) (35).

The phage genomes were analysed using the PHASTER bioinformatic tool, to search for filamentous phages (36). The sequences identified by PHASTER were confirmed manually by searching the disrupted tRNA sites (attB). The genes were annotated using the RAST server (37), HMMER (hmmer.com), Protein BLAST (38), and HHpred (39). The anti-phage defence systems were identified using the “The Procaykotic Antiviral Defense Locator (PADLOC)” tool (40).

### Phylogenetic and homology study

The genome sequences of the filamentous phages were aligned by the CLUSTAL method, and a maximum likelihood tree was constructed using Molecular Evolutionary Genetics Analysis (MEGA) software, version 11 (41).

Homologous analysis of the genomic sequences was done by Average Nucelotide Identity (ANI) with the ANI calculator tool (http://enve-omics.ce.gatech.edu/ani/) and by the BLAST Ring Generator Image (BRIG) (42). Finally, the homology of the protein sequences of the phages was determined using Easyfig 2.2.5 software (43).

### Phage propagation and purification

Cultures of *P. aeruginosa* PA01 or PA14, depending on the phage propagated, were grown overnight at 37°C and 180 rpm. The following day, the phage was propagated by the two-agar layer method (44). Briefly, the overnight culture was diluted 1:100 and incubated until the optical density at a wavelength of 600 nm (OD_600_) reached 0.5. An aliquot of 200 µl of the bacterial culture was mixed with 100 µl of the phage of interest. Soft TA (0.5% NaCl; 1% tryptone; 0.4% agar) was then added and the mixture was spread over a TA solid agar layer. The plates were incubated at 37°C for 24 h. The propagated phage was recovered by washing the plate with SM buffer (0.1 M NaCl, 1 mM MgSO4, 0.2 M Tris–HCl, pH 7.5); 1% chloroform was then added and the suspension was incubated for 20 min. Finally, the lysate was centrifuged and the supernatant containing the phages was recovered and stored at 4°C.

### Phage infection assays: spot test and infection curve

A spot test was conducted, as described by Kutter et al. (45), with minor modifications, to determine the sensitivity of the clinical isolates to the 4 lytic phages. Briefly, several plates were prepared by the double agar method with the host strain tested. Two µl of a suspension containing the phage of interest (10^9^ PFU/ml) was added to the top agar. The plate was incubated at 37°C for 24 h and the plate was analysed. Infection was considered positive (sensitive) if a clear or turbid spot was observed and negative (resistant) if no spot was observed. The tests were conducted in triplicate and were considered positive when all the replicates clearly showed a spot.

As some positive spots can occur as a result of abortive infection mechanisms or “lysis from without” and not from a productive infection (11), infection curve analysis in LB broth medium was conducted for the phage-host combinations that yielded positive spot tests. Infection curve analysis was conducted by combining 10^7^ CFU/ml of the clinical isolate selected and 10^8^ PFU/ml of the phage selected in 200 µl of LB broth in a 96 well microplate and incubating for 24 h at 37°C in an Biotek Epoch 2 (Agilent). Productive infection was assumed to have taken place when the OD was significantly lower than the control in the exponential phase.

### Relationship between defence systems in filamentous phages and host phage resistance

The probability of the host being resistant or sensitive when carrying a Pf-like phage was calculated as PR=P(R|Pf) or PS=P(S|Pf), where R is resistant and S is sensitive. The relationship between the presence of complete defence systems and the phage resistance of the host was also calculated as a differential probability for each defence system in those strains carrying a Pf-like phage, as dPR=P(DF|R)-P(DF|S), where DF is the defence system. Values of dPR close to 1 indicate overrepresentation of defence systems in resistant interactions, while negative values indicate overrepresentation of defence systems in sensitive strains (46).

## Supporting information

Fig S1. Infection curves for lytic phages and Pf-like carrier isolates yielding positive results in the spot test.

Fig S2. Infection curves of lytic phages and non-Pf-like carrier isolates yielding positive results in the spot test.

