## Supplementary figures and images for "«Involvement of Pf-like phages in resistance to phage infection in clinical isolates of *P. aeruginosa* from cystic fibrosis patients*»*"

### Fig S1. Infection curves for lytic phages and Pf-like carrier isolates yielding positive results in the spot test.

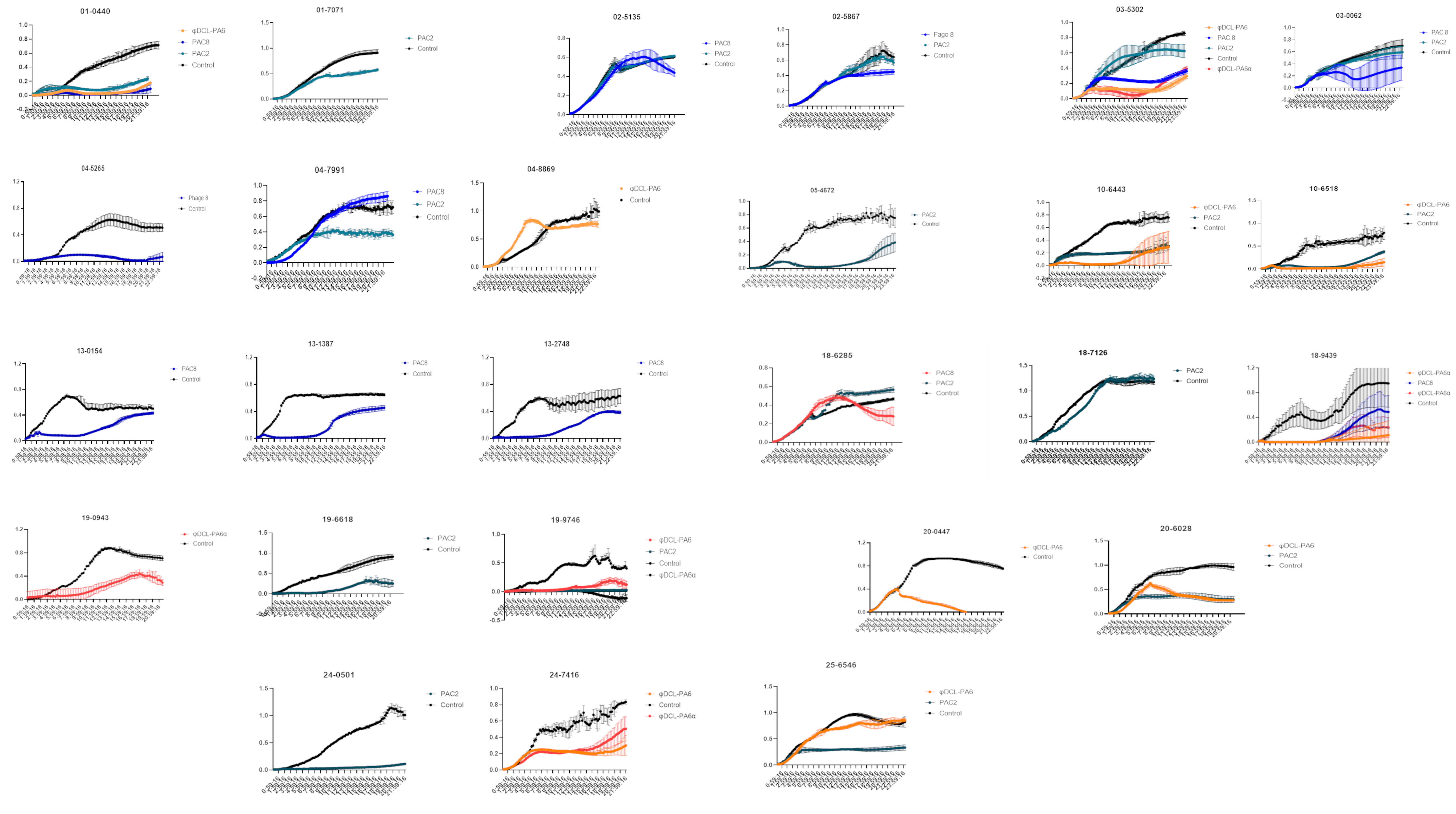

### Fig S2. Infection curves of lytic phages and non-Pf-like carrier isolates yielding positive results in the spot test.

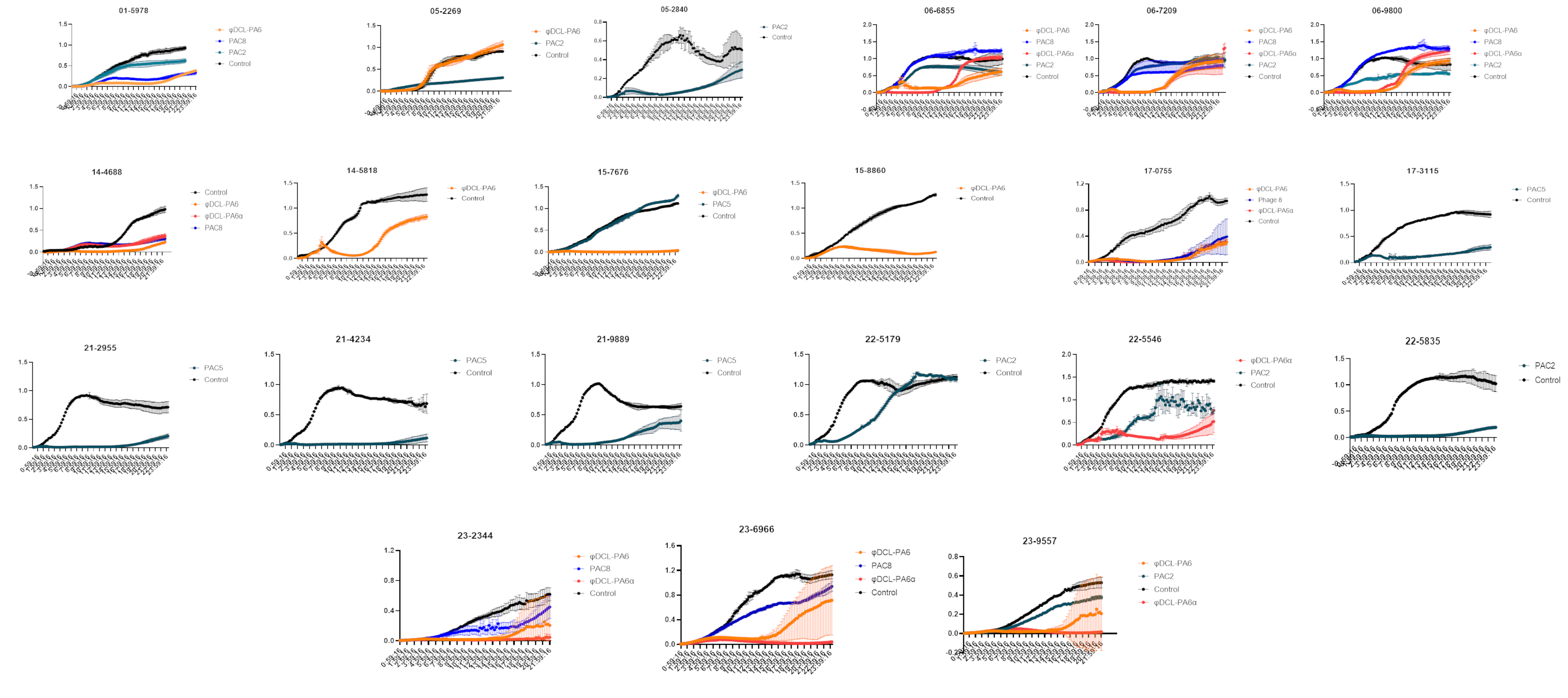
